# Priority effects and non-hierarchical competition shape species composition in a complex grassland community

**DOI:** 10.1101/253518

**Authors:** Lawrence H. Uricchio, S. Caroline Daws, Erin R. Spear, Erin A. Mordecai

## Abstract

Niche and fitness differences control the outcome of competition, but determining their relative importance in invaded communities – which may be far from equilibrium – remains a pressing concern. Moreover, it is unclear whether classic approaches for studying competition, which were developed predominantly for pairs of interacting species, will fully capture dynamics in complex species assemblages. We parameterized a population dynamic model using competition experiments of two native and three exotic species from a grassland community. We found evidence for minimal fitness differences or niche differences between the native species, leading to slow replacement dynamics and priority effects, but large fitness advantages allowed exotics to unconditionally invade natives. Priority effects driven by strong interspecific competition between exotic species drove single-species dominance by one of two exotic species in 80% of model outcomes, while a complex mixture of non-hierarchical competition and coexistence between native and exotic species occurred in the remaining 20%. Fungal infection, a commonly hypothesized coexistence mechanism, had weak fitness effects, and is unlikely to substantially affect coexistence. In contrast to previous work on pairwise outcomes in largely native-dominated communities, our work supports a role for nearly-neutral dynamics and priority effects as drivers of species composition in invaded communities.

## Introduction

Understanding the long-term outcome of competition in diverse species assemblages is a major goal of ecology, but measuring outcomes empirically is often infeasible when dynamics proceed slowly relative to the timescale of experiments. Theoretical coexistence models based on species differences offer a powerful alternative for predicting competitive outcomes (Burdon and Chilvers, 1974; Tilman, 1980; Holt et al., 1994). However, linking models to real ecological communities is challenging because realistic parameter estimates are rarely available for species interacting within a single ecological context. Moreover, existing theoretical approaches often make simplifying assumptions–such as reducing analyses to pairs of interacting species rather than considering the full dynamics of the assemblage–that may affect the inferred outcomes. As a result, considerable uncertainty remains about the relative importance of different proposed diversity-maintenance mechanisms, and the outcome of competition can be difficult to predict.

Among several existing theoretical frameworks for species interactions, modern coexistence theory clarifies the mechanisms through which species differences affect the outcome of competition via their effects on niche differences and fitness differences (Chesson, 2000). Niche differences are the differences between species in resource use, natural enemies, habitat requirements, or other factors that force percapita impacts of competition within species to exceed those between species (Chesson, 2000). These niche differences cause population growth to be negatively frequency-dependent because species achieve their highest per-capita growth rates when they are rare and interact with mostly heterospecific competitors, while per-capita growth declines as species become common and interact with many conspecific competitors. By contrast, in this framework, fitness differences are those differences between species that determine per-capita growth rates independent of relative abundance, often including differences in fecundity, resource acquisition, or survival. Coexistence occurs when niche differences, which generate negative frequency-dependent population growth or ‘stabilization’, are strong enough to overcome fitness differences between species (Chesson, 2000). Alternatively, priority effects can occur when population growth is positively frequency-dependent, allowing any species that is initially more common to exclude others but preventing species from invading when rare in the presence of competitors (Mordecai, 2011; Fukami et al., 2016). While assessing competitive outcomes in pairs of species with known parameter values is straightforward in this framework, predicting multispecies outcomes remains a challenge (Saavedra et al., 2017; Barabás et al., 2018).

Modern coexistence theory has motivated a suite of studies that empirically measure the strength of niche differences and fitness differences, and ultimately the outcome of competition, in natural plant or algal communities. Many of these studies have emphasized the influence of stabilizing niche differences on species coexistence (Turnbull et al., 2005; Adler et al., 2006; Levine and HilleRisLambers, 2009; Adler et al., 2010; Narwani et al., 2013; Godoy et al., 2014; Kraft et al., 2015; Godoy et al., 2017). However, almost all empirical applications of modern coexistence theory use pairwise growth rates when rare to assess community-wide outcomes (*e.g.,* Adler et al., 2007; Levine and HilleRisLambers, 2009; Adler et al., 2010; HilleRisLambers et al., 2012; Mordecai, 2013; Godoy et al., 2014; Kraft et al., 2015; Mordecai et al., 2015). A growing body of theory shows that outcomes of competition may depart from pairwise predictions when all species are analyzed simultaneously (*e.g.,* Case, 1995; Allesina and Levine, 2011; Saavedra et al., 2017; Mayfield and Stouffer, 2017), and that even two-species models can depart from the predictions of invasion analyses in some settings (Barabás et al., 2018). Deviations from pairwise predictions may originate from indirect effects on competitor density, non-hierarchical interactions, in which competitive impacts are intransitive, or higher-order interactions, in which the per-capita impact of one species on another is mediated by interactions with a third species. In complex communities it is theoretically possible to observe priority effects within groups of species with high niche overlap and fitness similarity, which occur alongside deterministic coexistence or competitive exclusion between groups of species with larger niche differences and fitness differences. Despite these theoretical developments, there are few empirical examples of communities in which multispecies outcomes depart from two-species predictions (but see Mayfield and Stouffer, 2017; Godoy et al., 2017), although alternative stable states have been empirically documented outside the coexistence literature (Staver et al., 2011; Carpenter et al., 2011; Fukami et al., 2005). It is unknown whether departures between pairwise and multispecies predictions are a common biological phenomenon or a quirk of mathematical models. Moreover, it is not clear how ecological contexts – such as the coevolutionary history of the competing species and the level of disturbance – may influence the likelihood of complex multispecies dynamics as compared to negative frequency-dependent stabilization (Ocampo-Ariza et al., 2018).

We hypothesized that complex multispecies dynamics would be likely to emerge in recently invaded systems, in which competing species have not had long coevolutionary histories and hence may not have had sufficient time to evolve into distinct niches (MacDougall et al., 2009). Such systems may be far from equilibrium, and may represent an opportunity to study complex multispecies dynamics in the absence of strong stabilizing mechanisms driven by niche differentiation. Moreover, predicting competitive outcomes in recently invaded systems is a pressing concern for conservation, and understanding the processes that have the largest impact on the persistence of native species could help preserve endemic biodiversity. In this study, we investigated competitive outcomes in California grasslands, one of the most ecologically important and widespread ecosystems in the Western U.S. and a key reservoir of endemic biodiversity (Myers et al., 2000). Simultaneously, California grasslands are heavily invaded and human-impacted since the 19th century (Mack, 1989); exotic annual grasses now dominate while native perennial bunchgrasses and annual forbs have declined to very low density throughout much of their extent. In the long term, it is not clear whether the exotic annual grasses will competitively exclude native perennial grasses, whether native grasses will stably persist (at reduced densities), or whether priority effects will determine whether native or exotic grasses dominate at a local scale.

Here, we combine field experiments with mathematical models to predict the outcome of competition in a community of five co-occurring native and exotic grasses in a grassland ecosystem. We develop and empirically parameterize population dynamics models for the species in this complex community. The models incorporate a suite of demographic rates experimentally measured for three different exotic annual grasses and two native perennial grasses at a single Northern California grassland site and include the impact of fungal pathogens on those demographic rates, which may affect competitive outcomes by altering fitness differences or by inducing negative frequency dependence of growth rates (Mordecai, 2011, 2013). We estimate parameters of the model statistically from field observations of seed output, germination, establishment, seed infection, and adult survival for each of five competing species of exotic annual and native perennial grasses. Using the parameterized model, we study the multispecies outcome of competition, with and without pathogens. Specifically, we ask: (1) How strong are intra- and interspecific competition between native and exotic grass species? (2) What is the predicted outcome of competition? (3) Do complex outcomes, including priority effects and/or intransitive competition, emerge from the dynamics of the full multispecies assemblage? (4) How strongly do foliar and seed pathogen impacts on demographic rates influence the outcome of competition?

## Methods

### Empirical data collection and parameter estimation

#### Study system

We chose five focal species that are highly abundant and widespread in California grasslands: exotic annual grasses *Avena barbata*, *Bromus diandrus*, and *Bromus hordeaceus*, and native perennial bunchgrasses *Stipa pulchra* and *Elymus glaucus*. For each species, we measured a full complement of demographic rates that determine population growth: seed germination, seedling establishment, the impact of competition on per-capita seed production, and over-summer survival (for both seedling and adult perennial bunchgrasses). We also assessed seed infection and survival with a common fungal pathogen that causes a disease called “black fingers of death,” which can kill native and exotic grass seeds (Beckstead et al., 2010). All experimental work was conducted in Jasper Ridge Biological Preserve, located in San Mateo County, California (37°24’N, 122°13’30”W; 66 - 207 m). This 485-hectare biological preserve is managed by Stanford University and has a Mediterranean climate with cool, wet winters and warm, dry summers (mean annual precipitation = 622.5 mm) (Ackerly et al., 2002). All demographic rates (except the perennial seedling to adult transition probability; see below) were measured in winter 2015 to spring 2016 (the 2016 growing season), an average rainfall year with 601 mm total precipitation distributed over 20 precipitation events.

#### Competition experiment & transect sampling

Seed output is an important component of plant reproduction, and is strongly impacted by competitor density for many plants (Mordecai, 2013; Mordecai et al., 2015). We measured seed output as a function of competitor density by varying the density of each plant species in monoculture and in mixed-species plots and measuring per-capita impact on seed production for each competitor species (Mordecai, 2013; Levine and HilleRisLambers, 2009). The 1-m^2^ competition plots were randomly assigned to five competitor density treatments ranging from 10% - 100% of each species’ estimated natural density in monoculture, for each of the seven background ‘species-groups’ (*A. barbata, B. diandrus, B. hordeaceus, E. glaucus* seedlings, *E. glaucus* adults, *S. pulchra* seedlings, and *S. pulchra* adults). We also cleared 4-m^2^ plots of competitors to measure seed output on individuals of each species at low competitor density (though natural recruitment from the seed bank meant that these plots were not completely clear of competition). We used weed matting, weeding, and seed addition to manipulate competitor densities. At the end of the 2016 growing season we measured seed output from up to three focal plants per species per plot (as available). See Supplementary Materials for a complete description of the competition experiment.

We also measured the impacts of competition on seed production in six transects that naturally varied in plant composition. Each transect varied in grass composition from native perennial-dominated at one end to exotic annual-dominated at the other, with five 1-m^2^ plots spanning each transect spaced approximately 5-10m apart. For each focal plant in the transects we estimated seed output, in both the 2015 and 2016 growing seasons (see Supplementary Materials).

To assess pathogen damage and its relationship with seed production and competitor density, we visually estimated leaf area damaged by fungal infection on a subset of leaves on each focal plant in both the transects and experimental plots. We used these data to quantify the impact of foliar fungal infection on seed output, as well as the impact of plant species local relative abundance on pathogen damage. We note that this approach could underestimate the impact of fungal infection if resistance is generally costly, and most surveyed plants were actively fighting infections.

Our attempt to manipulate competitor densities in the experimental plots was only partially successful because of recruitment from the seed bank. As a result, we combined all focal plants into a single data set of seed outputs and observed competitor densities (rather than using the target densities for the experimental plots). Our final dataset included 439 *A. barbata*, 399 *B. hordeaceus*, 319 *B. diandrus*, 387 *S. pulchra*, and 234 *E. glaucus* individuals (see Supplementary Materials for further details).

We used Markov Chain Monte Carlo (MCMC) to infer parameters relevant to seed output (*λ*), competition (*α*), and foliar infection (*β*) from our competition experiment data (see dynamic model section and Table 1 for parameter definitions), and performed simulation-based validation of our estimation procedure. Full details of this estimation procedure and its validation are provided in the Supplementary Materials, along with a more detailed description of the experimental design. We repeated the parameter inference procedures on subsets of the data that included either only the experimental plots or only the transects, and found that parameter values were of similar magnitude and with overlapping posterior distributions (see Fig. S7 and Supplementary Materials).

**Table 1:** Descriptions of parameters and variables that are relevant to our population dynamic model. Where not otherwise specified, *j* subscript refers to species *j*.

| Parameters & variables | Description |
| --- | --- |
| $\alpha_{jl}$ | per-capita competition impact of species $l$ on species $j$ |
| $\beta_j$ | impact of foliar fungal infection per unit area infected |
| $A_j$ | mean leaf area infected |
| $\lambda_j$ | mean seed output in the absence of infection and competition |
| $f_j(t)$ | compound impact of competition and infection on seed output |
| $S_j$ | seed output after accounting for competition and infection |
| $g_j$ | germination fraction |
| $\gamma_j$ | seed survival rate in presence of BFOD pathogen |
| $\nu_j$ | seedling survival rate |
| $\phi_j^I$ | establishment probability for infected seedlings |
| $\phi_j^U$ | establishment probability for un-infected seedlings |
| $D_l$ | density of competitor species $l$ per m<sup>2</sup> |
| $\nu_j^0$ | seedling transition rate in the absence of competition |
| $\nu_j(t)$ | seedling transition rate after accounting for competition |
| $\alpha_s$ | impact of seedlings on seedling-adult transition |
| $\alpha_a$ | impact of perennials on seedling-adult transition |
| $\alpha_n$ | impact of exotic annuals on seedling-adult transition |
| $D_s$ | density of seedlings per m<sup>2</sup> |
| $D_a$ | density of exotic annuals per m<sup>2</sup> |
| $D_n$ | density of perennials per m<sup>2</sup> |
| $P_j^I$ | proportion of seedlings infected during seedling-adult transition |
| $\xi_j$ | rate of perennial adult over-summer survival |
| $n_j^a(t)$ | number of annual seeds at time $t$ |
| $n_j^p(t)$ | number of perennial seeds at time $t$ |
| $a_j(t)$ | number of perennial adults at time $t$ |

#### Germination data

We planted marked seeds of each focal species to track germination and seedling establishment in 30 plots of 25 individuals of each species in November 2015. We recorded their status weekly for four weeks in January - February as missing (M), alive (A), or alive with *>* 50% of leaf area with pathogen damage (P). We used a Markov model to estimate the probability of establishing, surviving, and becoming heavily infected for each species by calculating weekly transition probabilities between these three states by plant species, then using the matrix to project forward four weeks to estimate the overall probability of germinating and establishing. We repeated this calculation both with pathogens present and with pathogens removed (*i.e.*, as if all individuals experienced the establishment rates of uninfected individuals), in order to understand the impact of pathogens on seedling establishment (Fig. S1 shows inferred germination *g*_*j*_ and establishment *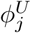* and *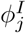* for uninfected and infected individuals, respectivelyfor each species *j*).

#### Seed survival and infection data

Seeds that remain dormant in the seed bank may contribute to population growth when they germinate in future years. To assess survival of non-germinating seeds between years, we buried 30 mesh seed bags contained five sewn compartments of 100 seeds of each focal plant species 2-5 inches deep from November 2015 - July 2016. We retrieved all intact seeds and scored them for “Black Fingers of Death” (BFOD) pathogen damage (Beckstead et al., 2010; Mordecai, 2013), which is likely caused by one of the numerous *Pyrenophora* spp. that are common in these sites (Spear and Mordecai, 2018). We tested all intact, non-germinated seeds for viability using germinatino, gibberellic acid, and cut tests as in previous work (Mordecai, 2012). We parameterize the fraction of seeds killed from BFOD infection as *γ*_*j*_ for species *j* (Fig. S2)

#### Adult survival data

To assess the survival of *S. pulchra* and *E. glaucus*, we marked 200 adult perennial bunchgrasses with flags in June 2015 at four sites for each species, and revisited prior to senescence in June 2016. We recovered 167 *S. pulchra* and 170 *E. glaucus* flags on the second visit and recorded them alive or dead based on observnig green tissue and/or new fruiting stalks. We fit binomially distributed models of survival probability for each species using beta-distributed priors with parameters *α* = 1, *β* = 1 (Fig. S3).

#### Perennial transition from seedling to adult

We modeled perennial seedling over-summer survival as a function of competitor density and per-capita competitive effects using data and fitted parameters from previously published work (Mordecai et al., 2015). Briefly, we modeled perennial seedling over-summer survival as a Beta-Binomially distributed process where for each observation *k*, the number of surviving seedlings maturing into adulthood was:

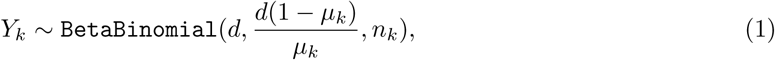

where *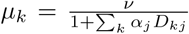* is the expected value, over all competitor species *j*. Here, *v* is the per-capita survival probability in the absence of competition, *α*_*j*_ are the per-capita competitive effects of individuals of species *j*, and *D*_*kj*_ are the number of individuals of species *j* in observation *k*. In the Beta-Binomial distribution *d* affects the mean and variance, and *n*_*k*_ is the number of trials in observation *k*. The equation for *Y*_*k*_ is parameterized so that its mean is equal to *µ*_*k*_. Since we lacked the data to separately estimate *α*_*j*_ for each individual species in our sample, we estimated an *α*_*j*_ for perennial adults, perennial seedlings, and exotic annuals, respectively.

### Population dynamic model

#### Model structure

We developed a dynamic model that captures the effects of competition and fungal infection on growth rates for each grass species in our system, which is structurally similar to models developed in previous work (Mordecai, 2013; Mordecai et al., 2015). All parameters and variables in the population dynamic model are listed in Table 1. To model the effects of competition on the population growth of each species, we suppose that each plant of species *j* has a seed output of *λ*_*j*_ in the absence of competition, and that competition and infection reduce seed output by

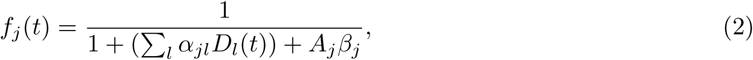

where *D*_*l*_(*t*) is the density of competitor species *l* per m^2^ at time *t* and *A*_*j*_ is the mean leaf area that was infected on individuals of species *j*. The expected seed output of a single individual *S*_*j*_(*t*) is then

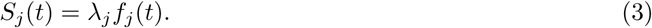

In the absence of transmisison information we do not explicitly model transmission of fungal pathogens, but simply assume that each individual is infected at the mean level of observed foliar fungal infection for species *j*.

For each species *j*, we suppose that a fraction *g*_*j*_ of the seeds germinate, and among these a fraction *γ*_*j*_ are infected with a fungal seed pathogen (BFOD). Seeds that germinate and are not infected may then establish, or may be eliminated before they ever produce seeds. We supposed that the probability of establishment depends on foliar fungal infection status, and separately estimated the probability of establishment for infected *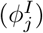* and uninfected *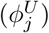* seedlings for each species *j*. The proportion of seedlings that are infected during the transition is given by *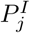*.

Perennial seedlings are presumed to be weak competitors and are not included in the competition model and produce no seeds in the first year of life (Mordecai et al., 2015), but are subject to competition in their survival over their first summer. Perennial seedlings survive at rate *v*_*j*_(*t*), given by

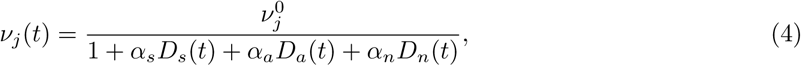

where *D* is the density of competitors, the subscripts *s*, *a*, and *n* represent perennial seedlings, perennial adults, and annuals, respectively, and *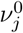* is the survival rate in the absence of competition. For this aspect of the model, we group all annual and perennial plants together instead of computing each species competitive effect separately because we lacked the data necessary to infer the parameters for each species individually. Perennial seedlings that survive the summer then become adults in the following year. Perennial adults are also subject to over-summer mortality at rate (1-*ξ*_*j*_), modeled as competition-independent because deaths of adult bunchgrasses are relatively rare (Fig. S3).

We model germination rate *G*_*j*_ after accounting for infection and establishment (see “Seed survival and infection data” section) with

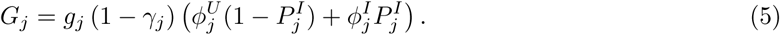

Additionally, we suppose that the proportion of seeds remaining in the seed bank after germination and infection *B*_*j*_ is given by

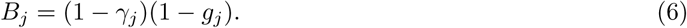

At each time step, for each species *j* we track the number of seeds *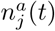* for exotic annuals, or the number of seeds *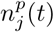* and adults *a*_*j*_(*t*) for perennials. Combining across seed production, competition, foliar fungal infection, fungal seed infection, germination, establishment, and over-summer survival, the full population growth equations are

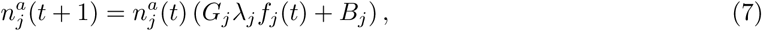

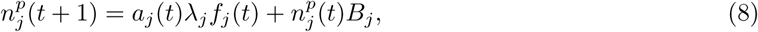

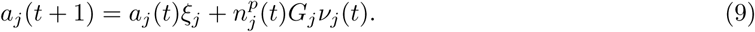

The germination, adult survival, seed infection, and seedling transition parameters were estimated as described in the previous sections. The competition (*α*), seed output (*λ*), and foliar infection burden (*β*) were estimated using custom MCMC software, as described in the supplemental methods. Simulations of each species in monoculture shows a strong positive correlation with observed monoculture densities in the field (Fig. S6), suggesting that our model parameterization with independently measured demographic rates accurately describes the system.

#### Separating infectious processes from competition

Initially we considered three broad classes of models that 1) did not include the impacts of infection, 2) included only foliar infection, and 3) included both foliar infection and BFOD pathogen. Since foliar infection had a negligible impact on dynamics and competitive outcomes, we compare models that include only foliar fungal infection to models that include both foliar infection and BFOD. The foliar infection model includes both the parameters relevant to infection of perennial seedlings during their transition to adults and infection of leaves of adult perennials and exotic annuals.

#### Inferring the outcome of competition

We used the parameterized models to investigate the outcome of competition using both pairwise and multispecies approaches, to assess any differences in the outcome across approaches and to understand potentially complex outcomes more mechanistically. First, we used the posterior parameter estimates and the population dynamic model to compute growth rate when rare (GRWR) for each pair of species. If a pair of species each have GRWR*>* 1 (i.e., log(GRWR)*>* 0, where log is the natural logarithm), then stable coexistence is predicted because the rarer species is expected to increase in prevalence, preventing either species from being excluded (Turelli 1978; but see Case 1995). Pairwise competition calculations were performed in the presence of fungal infection (*i.e.*, including all model parameters pertinent to foliar or seed fungal infection).

To calculate GRWR for each species pair, we sampled the relevant demographic rate parameters from their posterior distributions and simulated one species forward to its stable monoculture density, then computed the growth rate of an invader of another species. For perennials, we computed GRWR as the dominant eigenvalue of the 2x2 matrix describing the rate of transition between seed and adult stages, which accounts for their stage-structured life history (Mordecai, 2013). For exotic annuals, we calculated GRWR by simulating the addition of a single individual to the monoculture species and calculating the growth rate in the second generation. We also calculated percent difference in GRWR (pdGRWR) as *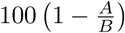*, where *B* is the GRWR in an empty plot, and *A* is the GRWR under invasion conditions.

Second, we investigated competitive outcomes for the full species assemblage using simulations. Because outcomes could be initial condition-dependent, we initialized our simulations at a range of observed densities from the transects. We simulated 600 years and report results of species mixtures at this time point (Mordecai, 2013). For species that were not present in a sampled transect, we introduced a pseudocount of one individual so that each species is initially present in each simulation. We consider species with population sizes exceeding one individual at the end of the simulation to have “persisted”, while species with less than one individual are “excluded.”

Lastly, we performed invasion experiments to better understand the dynamics underlying some of the coexistence outcomes that were predicted under our model. In these experiments, we compared the ability of *E. glaucus* to invade established *S. pulchra* plots with and without the presence of an exotic annual competitor. We considered only the subset of posterior parameter estimates in which *S. pulchra* was predicted to coexist with *B. diandrus*, excluding all replicates in which single species dominance was predicted. We simulated 100 years of *S. pulchra* growth in the absence of competitors to allow it to establish. Subsequently, at time *t* = 100, we either invaded *B. diandrus* or allowed *S. pulchra* to continue in the absence of competitors for another 100 years, and finally we introduced a single *E. glaucus* individual at time *t* = 200. We summarized these experiments by calculating the proportion of posterior samples in which *S. pulchra* is able to prevent *E. glaucus* from invading after 600 simulated years post-invasion. We sampled one set of parameters randomly from among the posterior estimates to display this invasion experiment, but note that we vary the timing of the invasions for the purpose of visualization (Fig. 4).

**Figure 4:**
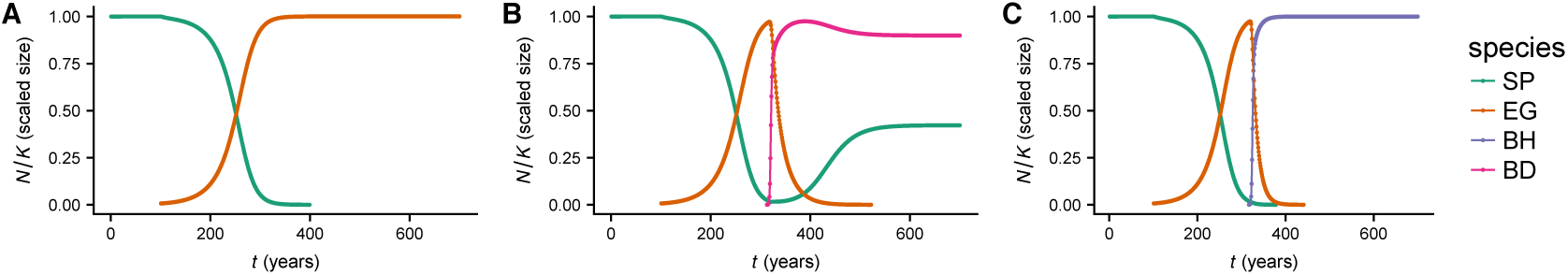
Effect of exotic annuals on invasion dynamics. A-C correspond to a single set of parameter estimates sampled from the posterior distribution. A: *EG* invades at *t* = 100, and is able to establish and eventually replace *SP*. B: *EG* invades at *t* = 100, and is able to establish and gradually replace *SP*. *BD* invades at *t* = 220, and coexists with *SP*, reversing the invasion of *EG* and preventing *EG* and *BH* from invading in the future. C: *EG* invades at *t* = 100, and is able to establish and gradually replace *SP*. *BH* invades at *t* = 220, and out-competes both perennials. *BD* cannot invade due to the priority effect between *BD* and *BH*. Units on the y-axis correspond to population sizes relative to monoculture density (Abbreviations: perennial native species (*S. pulchra* (SP), *E. glaucus* (EG)), annual exotic speices (*B. hordeaceus* (BH), and *B. diandrus* (BD))).

## Results

### Evidence for strong interspecies competition

We calculated the pairwise log(GRWR), where zero is the threshold for invasion, of each species invading a monoculture of each other species to assess the pairwise outcomes of competition (Fig. 1A). We also calculated the proportional decrease in GRWR relative to the growth rate in the absence of competition to assess the overall impact of competition on population growth (pdGRWR, Fig. 1B). The values plotted in Fig. 1 represent the mean over 250 independent samples of the parameters from their posterior distribution. If there were strong niche differences between all pairs of species, each species would strongly constrain its own growth rate but to have a relatively weak competitive impact on other species. By contrast, we observed high niche overlap in this system, in which interspecific competition constrains invasion growth rates as much or more than intraspecific competition (Fig. 1A). Along the diagonal, where each species competes with itself at its stable monoculture density, the log(GRWR) was always very near zero (Fig. 1A). When the native perennial species (*E. glaucus* and *S. pulchra*) were the invaders, they were constrained to log growth rates close to zero by all competitors except *A. barbata*, which is unable to constrain either species. By contrast, neither perennial species reduced the log(GRWR) of any of the annual species to below zero. *B. hordeaceus* and *B. diandrus* tightly constrained each others’ growth rates, but were not strongly constrained by the perennial competitors. *A. barbata* was a poor competitor in this community, and only constrained itself while being tightly constrained by both of its annual competitors.

**Figure 1:**
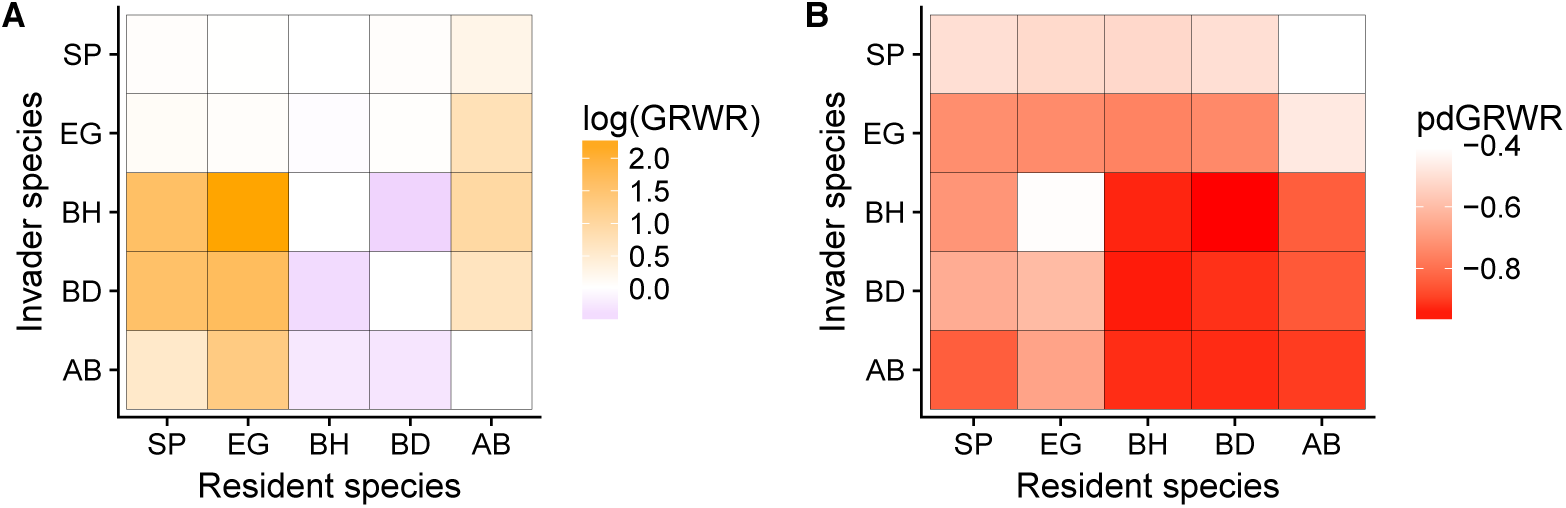
Strength of pairwise competition inferred from the growth rates of invaders introduced at low density to established monocultures of residents. A: log of growth rate when rare (GRWR) – log(GRWR)*>* 0 is the invasion criterion. Larger positive values (darker yellow/orange) represent high niche overlap, while darker purple represents less overlap. B: shows the proportional difference in each species growth rate in the presence of the resident species as compared to its growth rate in the absence of competition. p. (Abbreviations: perennial native species (*S. pulchra* (SP), *E. glaucus* (EG)), annual exotic species (*A. barbata* (AB), *B. hordeaceus* (BH), and *B. diandrus* (BD))).

These results are not driven solely by some species having low overall (competition-independent) growth rates, because competition reduced GRWR for nearly all species pairs (Fig. 1B). The strongest impacts of competition were of *B. diandrus* and *B. hordeaceus* on each other, resulting in growth rates reduced by 95 *-* 96.4% relative to the absence of competition. *E. glaucus* and *S. pulchra* were also subject to competition with each other, reducing *S. pulchra*’s growth rate 49.3 *-* 51% and *E. glaucus*’s growth rate by 72.7 *-* 73.6% relative to growth rates in absence of competition. The impacts of *B. diandrus* and *B. hordeaceus* on the perennials were of similar magnitude to the observed perennial-perennial impacts.

### Predicted community composition is exotic annual-dominated

We next sought to investigate competitive outcomes when all five species compete simultaneously. Although it would be possible to perform a multispecies invasion analysis, such analyses can provide misleading results because GRWR exceeding 1 does not necessarily imply stable persistence when more than two species are included (Case, 1995; Dormann and Roxburgh, 2005; Saavedra et al., 2017). Instead, we examined the outcomes of competition by directly assessing the final simulated community composition across a range of empirically observed densities as initial conditions. For this analysis (Fig. 2A) we included the impact of foliar fungal infection on demographic rates, but not the pathogen BFOD.

**Figure 2:**
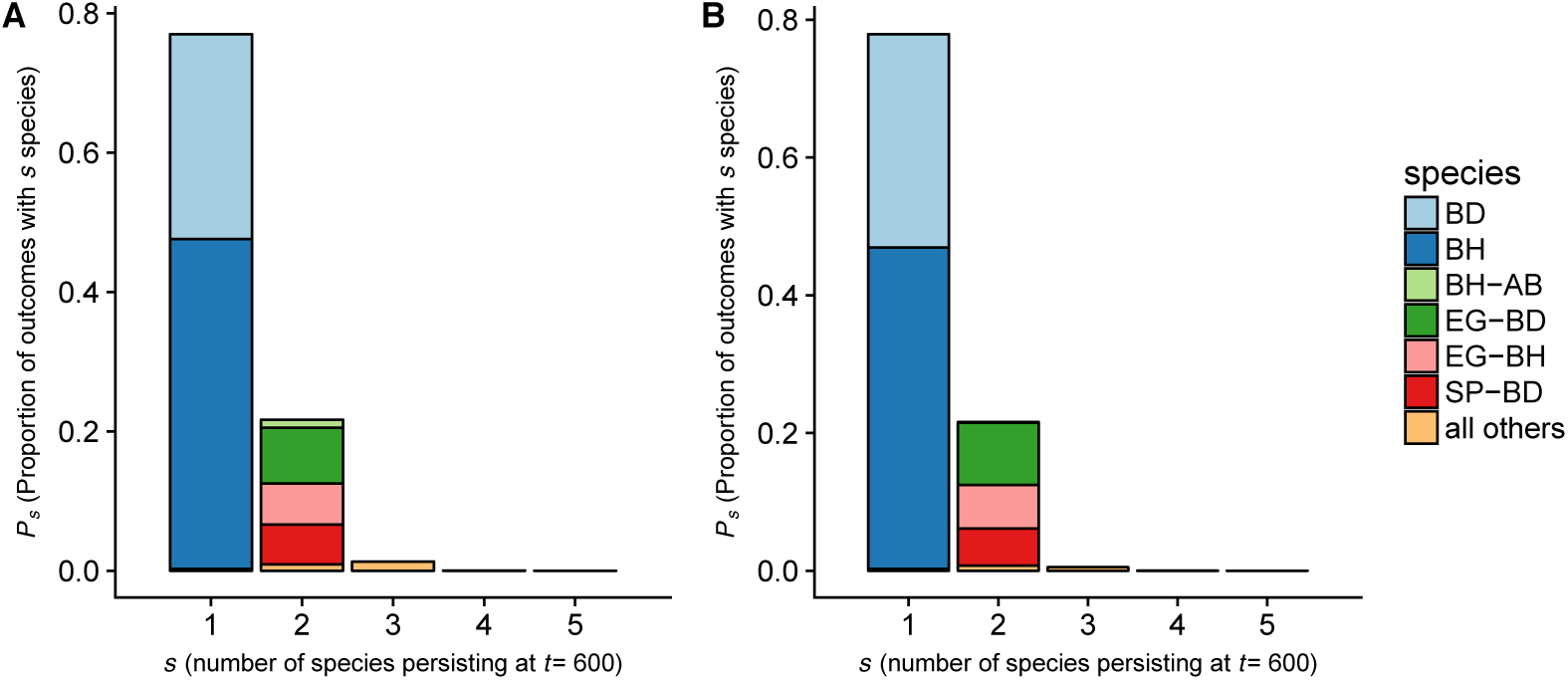
The proportion of outcomes in which a given species or species combination persisted after 600 generations in the population dynamic model. Each replicate represents a single sample from the posterior distribution of the model parameters, and initial competitor densities from a randomly sampled transect from among the set of transects that we measured. The x-axis corresponds to the number of species (*s*) that persisted over 600 simulated years, while the y-axis corresponds to the proportion of replicates (*P*_*s*_) that resulted in this outcome. Each bar is subdivided into the specific set of species that were observed to have log(GRWR)*>* 0 for a particular parameter set and transect sample. All compositions that occurred in under 1% of replicates are grouped into a single color. A: results from a model that includes foliar fungal infection but not BFOD; B: results from the model that includes both foliar fungal infection and BFOD. (Abbreviations: perennial native species (*S. pulchra* (SP), *E. glaucus* (EG)), annual exotic species (*A. barbata* (AB), *B. hordeaceus* (BH), and *B. diandrus* (BD))).

Based on our two-species competition observations (Fig. 1), we expected that either *B. hordeaceus* or *B. diandrus* would persist in most simulations because they have large growth rates even when invading a monoculture of native perennial competitors. Since this species pair displayed large interspecific competition coefficients, we also expected that they would rarely coexist. The pairwise invasion analysis also suggests that *A. barbata* should rarely persist given its low growth rate in the presence of its competitors, and that the native perennials *S. pulchra* and *E. glaucus* would sometimes persist, given that their mean replacement rates were typically near one (Fig. 1A). Consistent with these expectations, either *B. hordeaceus* or *B. diandrus* persisted in 99.7% of simulation replicates (Fig. 2A), while the two species almost never coexisted with each other (0.32% of replicates). *A. barbata* was nearly always excluded. Native perennials persisted in 22.0% of replicates, usually by coexisting with either *B. hordeaceus* or *B. diandrus*, but rarely coexisting with each other (0.39% of replicates).

The wide range of possible outcomes could reflect either uncertainty in the parameter estimates or dependence on the initial conditions of our simulations. To differentiate between these possibilities, we investigated persistence as a function of initial competitor densities (Fig. 3). If each species’ success is strongly dependent on initial conditions, this suggests that priority effects play a much larger role in determining community composition than uncertainty in parameter estimates, and vice versa. We observed that the perennial species’ persistence in multispecies competition was weakly dependent on initial community composition: the probability of *S. pulchra* persistence increased modestly with its initial density. By contrast, the annual species *B. hordeaceus* and *B. diandrus* showed striking priority effects driven by strong interspecific competition across the densities of all competitors except *A. barbata*, which had only modest effects on any species.

**Figure 3:**
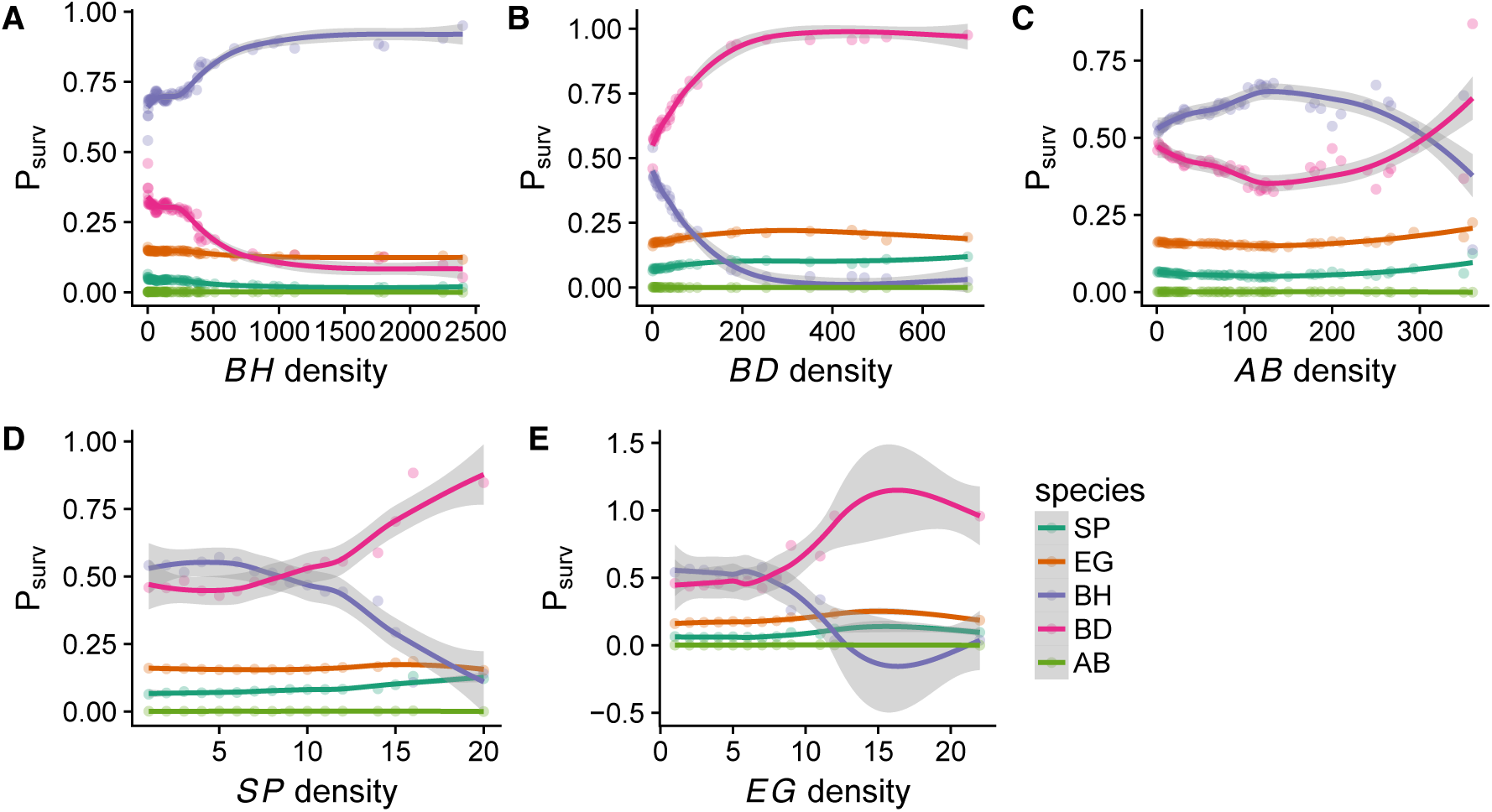
Effect of initial conditions on the outcome of competition. Estimated probability of persistence (*P*_surv_) of each species (colored points and lines) as a function of initial density of each species in the simulation (x-axis). Points represent the proportion of simulations resulting in persistence that were initialized at the corresponding density, while loess curves (and gray uncertainty envelopes) were fit with the function geom smooth in ggplot. (Abbreviations: perennial native species (*S. pulchra* (SP), *E. glaucus* (EG)), annual exotic species (*A. barbata* (AB), *B. hordeaceus* (BH), and *B. diandrus* (BD))). Priority effects occur when the probability of persistence depends strongly on initial density.

### Non-hierarchical competition between natives and exotics

Notably, the two native perennial species were not equally likely to coexist with each of the dominant exotic annuals: *S. pulchra* was much more likely to coexist with *B. diandrus* than *B. hordeaceus*. To better understand the dynamics driving these outcomes, we compared invasion simulations that included only the native perennials to simulations in which an exotic annual invader preceded the native perennial invader.

When the native perennials competed in the absence of exotic annuals, their dynamics are nearlyneutral, as indicated by the near equality intraspecific and interspecific competition strength (Fig. 1A). Although neutral dynamics cannot result in long-term stable coexistence, they are expected to result in relatively slow loss of one of the species (Adler et al., 2007). Nearly-neutral dynamics could result in priority effects if interspecific competition is slightly stronger than intraspecific competition. Consistent with these expectations, we observed slow replacement dynamics and priority effects when the native perennial species competed directly (Fig. 4A). When *S. pulchra* was introduced first, 36.4% of simulation replicates resulted in dominance by *S. pulchra*, whereas 99.9% of replicates in which *E. glaucus* was introduced first resulted in dominance by *E. glaucus*. The two perennial species coexisted in only 0.1% of replicates.

We next tested how the invasion of exotic annuals might alter competition outcomes between the two perennial species. When *B. diandrus* invaded simulated monocultures of *S. pulchra*, the priority effect preventing *E. glaucus* from invading was substantially stronger. Among replicates in which *S. pulchra* and *B. diandrus* coexisted, *E. glaucus* was able to invade in only 4.5% (as compared to 52.9% of these same parameter sets when *B. diandrus* was not included in the simulation). Hence, when coexistence was predicted between *S. pulchra* and *B. diandrus*, the presence of the exotic annual *B. diandrus* increased the likelihood of persistence for *S. pulchra* at the expense of *E. glaucus* (Fig. 4). These results suggest a strong impact of non-hierarchical competition on outcomes in this system, in which no single species can dominate and coexistence is only possible given specific arrival orders. These dynamics result in deterministic competitive exclusion or coexistence mediated by priority effects that make the final composition dependent on stochastic arrival order.

### Seed infection hampers a weak competitor, *A. barbata*

While the simulation and invasion results suggest that exotic annuals will dominate perennials and either competitively exclude them or reduce their population sizes, pathogens could reduce this effect if exotic annuals incur a higher fitness cost than perennials from infection. We repeated the simulation experiments in the previous section with the impact of fungal infection by BFOD included (Fig. 2B). BFOD decreased the overall success of *A. barbata*, which was already the weakest annual competitor. *A. barbata* persisted in 1.98% of replicates that excluded BFOD, but almost never persisted in its presence (0.12% of replicates). Species other than *A. barbata* were only minimally affected by BFOD, consistent with low rates of infection that we observed empirically (Fig. S2).

## Discussion

The long-term outcome of competition is difficult to observe directly in nature, so empirically parameterized mathematical models are critical for predicting when competitive exclusion, coexistence, or priority effects are most likely. To date, most studies have focused on pairwise outcomes in multispecies communities, emphasizing the relative strength of niche differences versus fitness differences (Levine and HilleRisLambers, 2009; Adler et al., 2010; Narwani et al., 2013; Godoy and Levine, 2014; Godoy et al., 2014; Kraft et al., 2015). By contrast, we found a large degree of niche overlap in an empirically parameterized model of an invaded California grassland (Fig. 1A). The model suggests that competition in multispecies assemblages can lead to complex outcomes that mix deterministic competitive exclusion or coexistence with stochastic priority effects that determine the identity of persisting species. Although a previous model predicted exotic annual dominance over native perennials, with a small probability of native coexistence with exotics, it did not predict priority effects among pairs of exotic or native species because it relied solely on a pairwise approach and composite parameter estimates from a range of species (Mordecai et al., 2015). Our work supports the results from a growing body of theoretical studies suggesting that competitive outcomes in complex communities can depart from predictions from pairwise species comparisons (Case, 1995; Allesina and Levine, 2011; Saavedra et al., 2017; Mayfield and Stouffer, 2017), and provides evidence that slowly occurring, nearly-neutral dynamics may play an important role in competitive outcomes (Adler et al., 2007). The results suggest that methods for assessing the outcome of competition in multispecies assemblages that move beyond invasion analyses are an important avenue for theoretical and empirical development (Saavedra et al., 2017; Barabás et al., 2018).

We found that the population growth model parameterized with field estimates of demographic rates accurately predicted species densities in monoculture (Fig. S6), suggesting that it can realistically capture equilibrium outcomes. In this five-species assemblage we found species pairs with high niche overlap—exotic annuals *B. diandrus* and *B. hordeaceus*; and native perennials *E. glaucus* and *S. pulchra*—and non-hierarchical competition such that no single species was dominant over all others (Fig. 1). This results in a complex posterior distribution of competitive outcomes that includes competitive exclusion of all other species by either *B. diandrus* or *B. hordeaceus*, or coexistence of one of these exotic annuals with one native perennial, either *E. glaucus* or *S. pulchra* (Fig. 2). The outcome of competition between the two congeneric exotic annuals was driven by strong interspecific competition, resulting in priority effects (Fig. 3). Among the subset of outcomes in which coexistence was predicted between *S. pulchra* and *B. diandrus*, the presence of *B. diandrus* usually prevented the subsequent invasion of *E. glaucus*, augmenting the priority effect exerted by *S. pulchra* on its native perennial competitor through non-hierarchical competition. Hence, invasion by an exotic species with moderate niche overlap may increase resilience to invasion by native competitors with high niche overlap. Since the two native species are predicted to compete nearly-neutrally in the model, almost all predicted outcomes precluded their long-term stable coexistence, meaning that *S. pulchra* may indirectly benefit from invasion by *B. diandrus* via its negative effect on *E. glaucus* (Fig. 4).

Studies applying modern coexistence theory have often documented substantial niche differences between species pairs, while variation in the magnitude of fitness difference determines the potential for coexistence or competitive exclusion. However, these studies typically explore communities in which species have long coevolutionary histories and belong to similar functional guilds (*e.g.*, native annual forbs) (Levine and HilleRisLambers 2009; Adler et al. 2010; Godoy et al. 2014; Kraft et al. 2015; Narwani et al. 2013; but see Godoy and Levine 2014). In such systems, we might expect that co-occurrence over long timescales has led species to minimize their fitness differences, maximize their niche differences, or both, leading to long-term stable coexistence. In invaded systems, exotic species must have fitness advantages, niche differences, or both with respect to the native species in order to have successfully invaded (MacDougall et al., 2009). Our study found support for large fitness advantages of invading species and modest niche overlap that was potentially small enough to promote coexistence (20% posterior probability). This is consistent with the possibility that competitive exclusion and priority effects may be common long-term outcomes in invaded communities. Despite the *a priori* expectation of strong stabilization within coevolved communities, our study did not identify substantial niche differences between the two native species or the two exotic *Bromus* species with an overlapping European home range. Instead, the results suggest that nearly-neutral dynamics determine competitive outcomes in some pairs of species with long coevolutionary histories.

Among the mechanisms that may maintain diversity in complex communities, intransitive competition has received a great deal of interest (Laird and Schamp, 2006; Allesina and Levine, 2011; Soliveres et al., 2015; Matías et al., 2018; Stouffer et al., 2018). However, a recent study using empirically parameterized population dynamic models found that intransitive competition was uncommon and not likely to promote coexistence among a set of 18 competing annual grass species (Godoy et al., 2017). In our study, although we found evidence for non-hierarchical competition, simulations suggested that it was unlikely to promote coexistence of more than two species. However, this does not imply that non-hierarchical competition has no role in diversity maintenance. Priority effects, like those reported here, could lead to a spatial patchwork of competitors, with disturbance and stochastic recolonization determining the distribution of competing species.

Previous empirical work on the outcome of competition in invaded California grasslands has suggested a wide range of possible outcomes and coexistence mechanisms, including (1) the requirement for previous disturbance to explain the invasion of exotics in perennial-dominated plots (Stromberg and Griffin, 1996; Seabloom et al., 2003; Corbin and D’Antonio, 2004), (2) competitive dominance of exotic annuals over *S. pulchra* (Dyer and Rice, 1997), and (3) spatial variation in outcomes determined by seed dispersal (DiVittorio et al., 2007) and habitat (Everard et al., 2009). A limitation of most previous work considering more than two competing species is that the empirical results were not linked to a dynamic model, making it difficult to project dynamics over long timescales. Our parameterized model is inconsistent with (1) because it predicts that exotic species can invade undisturbed perennial plots, and consistent with (2) because a portion of predicted outcomes led to unconditional exotic dominance, as predicted in a previous two-species model (Mordecai et al., 2015).

While our study did not attempt to assess environmental and spatial variation in competitive outcomes (*i.e.*, prediction (3) above), coexistence mechanisms occurring over broad temporal and spatial scales may be an important contributor to observed species distributions. The five focal species in our system have co-occurred at the study site and other California grassland sites for over a hundred years, implying that co-occurrence may be driven by non-equilibrium dynamics that play out over long timescales, or that stabilizing processes occur over larger spatial or temporal scales. Although soil moisture and nutrient availability vary spatially at both neighborhood and regional scales across California grasslands and precipitation varies several-fold between years, demographic studies of species interactions are regularly conducted at small spatial and temporal scales, as in the present study (Seabloom et al., 2003; Corbin and D’Antonio, 2004; DiVittorio et al., 2007). Environmental variation combined with the long lifespans of perennial bunchgrasses and the high seed production of annual grasses could provide ample opportunity for environmental fluctuations to promote coexistence over space or time (Warner and Chesson, 1985; Cáceres, 1997; Chesson, 2000; Chesson et al., 2005; Li and Chesson, 2018; Usinowicz et al., 2017). An alternative to larger-scale stabilizing processes is that priority effects occur at small spatial scales, while the turnover and random colonization of sites allows patches of different species composition to persist in a matrix of mutually non-invasible patches (Vannette and Fukami, 2014). Our results are consistent with this priority effect mechanism, and do not preclude the possibility of coexistence mechanisms occurring over larger scales. A major challenge for future work in systems with large and variable geographic ranges is to incorporate multi-scale spatial and temporal variation that may contribute to patterns of co-occurrence.

In addition to the outcome of competition, identifying explanatory mechanisms is paramount for understanding the assembly of ecological communities and their potential to respond to species invasions. Pathogens are widely hypothesized to promote species diversity by specializing on common host species, causing a demographic disadvantage to common species and an advantage to rare species (Mordecai, 2011, 2013; Bagchi et al., 2014; Petermann et al., 2008; Gilbert et al., 1994; Augspurger and Kelly, 1984; Augspurger, 1983). As with strong niche differences more generally, this Janzen-Connell effect has been demonstrated mostly in native-dominated plant communities (Augspurger, 1983, 1984; Augspurger and Kelly, 1984; Gilbert et al., 1994; Petermann et al., 2008; Bagchi et al., 2014). By contrast, pathogen impacts in this invaded grassland community differ in two important ways. First, the foliar fungal pathogens in this community typically infect multiple grass hosts, including native and exotic species (Spear and Mordecai, 2018). Second, the demographic impacts were mostly minimal, except the negative impact exerted by the Black Fingers of Death pathogen on *A. barbata* (Fig. 2B). Another study in this system found no evidence that pathogen load or impacts increased on locally common species, suggesting that coexistence is not substantially affected by fungal infection (Spear and Mordecai, 2018). However, because the two native perennial species compete almost neutrally in our model, even modest pathogen impacts could have a substantial effect; because their fitness differences are small, our study may have been underpowered to investigate this specific outcome.

Though the outcome of competition in invaded systems is often poorly understood, our work shows that stabilizing niche differences, and particularly the role of pathogens in that stabilization, are not ubiquitous at local scales in nature, especially in systems with short coevolutionary histories between species. Assessing the strength of stabilization and the outcome of competition is important not only for fundamental ecological understanding but also for maintaining resilient ecosystems as they respond to global change. Future work in other systems that are potentially far from equilibrium, such as invaded ecosystems, is necessary to test the generality of our results, and to predict the long-term outcome of competition in nature.

## Acknowledgments

We thank our editors and anonymous reviewers at *The American Naturalist* for comments that improved the quality of the manuscript. We acknowledge generous support and assistance from Teri Barry, Nona Chiariello, Phillippe Cohen, Joe Sertich, Reuben Brandt, Johannah Farner, Steve Gomez, Bill Gomez, Stuart Koretz, Cary Tronson, Divya Ramani, Ryan Tabibi, Esther Liu, Sandya Kalavacherla, Vidya Raghvendra, Jason Zhou, Virginia Parra, Claudia Amadeo-Luyt, Gaurika Duvar, Elizabeth Wallace, the Jasper Ridge Kennedy endowment fund, the Stanford University Vice Provost for Undergraduate Education summer research fellowship for undergraduates, and the Stanford University Raising Interest in Science and Engineering (RISE) summer internship program. We thank members of the Fukami, Peay, and Mordecai labs for helpful comments on the project, and Rodrigo R. Granjel, Ignasi Bartomeus, Oscar Godoy, and members of the Biotic Interactions and Global Change group for their preprint peer-review of our manuscript. LHU was supported by the NIH IRACDA fellowship (NIGMS grant K12GM088033). EAM was supported by a grant from the National Science Foundation (DEB-1518681).

## Supplemental methods & figures

### Competition experiment

To measure the impact of intra- and interspecific competition on seed production, we set up an experiment to vary the density of each plant species in monoculture and measured its per-capita impact on seed production for each competitor species (Mordecai, 2013; Levine and HilleRisLambers, 2009). We attempted to limit the seed bank by preventing seed set the year prior to the experiment. In the spring of 2015 before seed set, we laid down weed matting across a 35m x 35m area near the Sun Field Station in Jasper Ridge Biological Preserve. We cut holes in the matting to allow existing adult perennial plants (mostly *S. pulchra*) to survive. In fall 2015 before the first major rains, we removed the weed matting and established 210 1-m^2^ competition plots and 30 4-m^2^ competition-free plots. All plots had a 1-m untreated buffer area from all other plots on all sides.

We manipulated background competitor species density and fungal abundance in a factorial design. The competition plots were randomly assigned to treatments across five densities, seven background “species-groups” (*Avena barbata, Bromus diandrus, Bromus hordeaceus, Elymus glaucus* seedlings, *E, glaucus* adults, *Stipa pulchra* seedlings, and *S. pulchra* adults), and three fungal manipulations (fungicide, water controls, or fungal inoculation with liquid inoculum). Because the fungal inoculum was not successful in increasing fungal infection and fungicide only modestly reduced fungal load, we pooled all three treatments into a single dataset, and we do not describe fungal growth, inoculation, and fungicide methods here. In total there were two replicate plots of each background species x density x fungal treatment. We additionally established competition-free plots that aimed to have a single individual of each of the seven focal species-groups growing in the absence of competition with all other plants removed from the 2m x 2m plot area.

In the competition plots we sought to create a gradient of densities with each species-group as the ‘background’ species that comprised the dominant competitor environment, plus each remaining species-group present in low density to gauge the impact of the competitor environment on per-capita seed production for each species. Background competitor densities aimed for 10%, 20%, 40%, 80%, and 100% of the density of each plant species in monoculture, where monoculture densities were estimated by counting plants m^*-*2^ in the surrounding area. Monoculture densities were estimated at 10 adults m^*-*2^ for each of the two perennial species, 1,715 seeds m^*-*2^ for *A. barbata*, 3,209 seeds m^*-*2^ for *B. diandrus*, 12,544 seeds m^*-*2^ for *B. hordeaceus*, 4,585 seeds m^*-*2^ for *E. glaucus*, and 1,466 seeds m^*-*2^ for *S. pulchra*. In November, 2015, we added seeds and transplanted adult *S. pulchra* plants to achieve the desired density treatment in each plot, removing excess bunchgrasses from plots as needed. We transplanted adult *E. glaucus* plants from an area 1 km away into the plots in December, 2015. All transplants were watered to settle the soil. We then added approximately 10 seeds of each non-background species in January, 2016, marked with plastic cutlery stuck into the ground, and one adult of each perennial species, marked with a small plastic ring, to each plot to ensure that all focal species were represented in each plot. These would become the focal individuals on which we would assess seed production. In the competition-free plots we seeded approximately 10 seeds/species into separate locations within the 2m x 2m plots, each marked by plastic cutlery, plus one individual adult of each perennial species, marked by plastic rings. Finally, we reseeded 18 of the 240 plots with 75-100% of their original treatment seed weight in January, 2016 following low germination.

In addition to applying weed matting to prevent seed set the prior year, we weeded the plots extensively to remove target and non-target species and to thin plots to the desired density. However, despite these efforts a substantial density of non-target plants naturally recruited into the plots. As a result, although the experiment manipulated competitor density, it did not achieve the predetermined plant density targets. For this reason, we censused plant density and species at peak flowering in the spring, and used these actual competitor density estimates as inputs into all competition models.

We attempted to harvest seeds from all marked focal individuals in all plots in May-June 2016 when they matured but before they dehisced and dispersed. Because the seeds of *A. barbata* and *S. pulchra* dehisced very quickly at maturity and the timing varied within an individual, we harvested and counted the glumes of those individuals and estimated the number of seeds per glume to get an overall estimate of seed production (two seeds per glume for *A. barbata* and one seed per glume for *S. pulchra*).

### Transect plots

We also measured the impacts of competition on seed production in plots that naturally varied in plant composition, to understand the strength of competition in established plant communities. In the 2015 growing season, we set up six transects in the area surrounding the Sun Field Station at Jasper Ridge Biological Preserve. Each transect varied in grass composition from native perennial-dominated at one end to exotic annual-dominated at the other, with five 1-m^*-*2^ plots spanning each transect spaced approximately 5-10m apart. Individual plots were placed based on desired plant community compositions, thus plots were unevenly spaced (3 to 18m apart). Transects ranged from 20m to 50m in length. Three of the transects were dominated by *S. pulchra* at the perennial end, and the other three were dominated by *E. glaucus*. We surveyed plant density and community composition in the 2015 growing season. In 2016 we returned to these transects and used only the first, third, and fifth plot in each transect, representing perennial-dominated, intermediate, and annual-dominated plots, respectively. We paired each plot with two additional 1-m^2^ plots next to each original plot that were visually similar in composition, for a total of three sets of triplet plots per transect. In each triplet of plots, one received fungicide, one received a water control, and one received fungal inoculum. Again, the fungal inoculation was not successful at increasing pathogen infection, and fungicide only modestly decreased pathogen load, so we combined data from all treatments. We seeded approximately 20 seeds of each focal species, and transplanted an adult *S. pulchra* and/or an adult *E. glaucus* plant into each plot that lacked these species (seeding and transplanting were done on the same days as the competition plots described above). We censused the density of each plant species during peak flowering (when grasses are easiest to identify), then harvested all seeds from up to three individuals of each focal species present in each plot. We used data on the number of seeds per individual and competitor density to parameterize competition models, as described below.

### Competition & infection model

We use a simple model of competition between individuals on a patch, in which seed output *S* is affected by the density of competing individuals and infection burden. Since we observe that *S* has an over-dispersed distribution that is well described by a negative binomial (Fig. S4), we allow *S* to follow a negative binomial distribution.

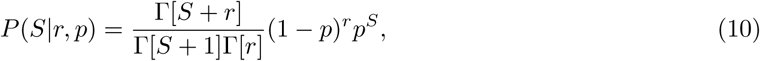

where Γ is the gamma function, and *p* and *r* are parameters that govern the mean and variance of the negative binomial distribution. We suppose that the expected fitness *f* of individual *i* in species *j* is a function of the number of individuals competing in the patch, and the amount of infected leaf tissue of each individual such that

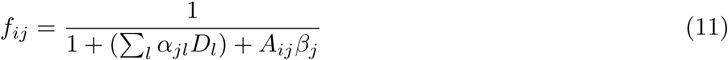

where *D*_*l*_ is the density of individuals of species *l*, and *α*_*jl*_ captures the strength of the impact of individuals of species *l* on *j* (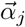 is the vector of *α*_*jl*_ values). *A*_*ij*_ is the area of infected leaf tissue of individual *i*, and *β*_*j*_ is the impact of foliar fungal infection per unit leaf area infected on fitness for individuals of species *j*. We constrain the *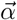* and *β* values to be non-negative, which captures our strong prior expectation that interactions between species and infection should have a deleterious effect on fitness. This competition model was previously studied in other systems (Mayfield and Stouffer, 2017). We incorporate the competition model into seed output *S* by allowing the mean of the negative binomial distribution to follow eqn. 11, while preserving the shape of the negative binomial. To do so, we allow the *r* parameter of the negative binomial to be given by

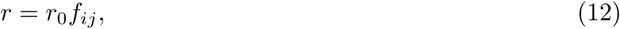

where *r*_0_ is the *r* parameter in the absence of competition and infection. This model formulation constrains the mean seed output to be decreased proportional to the plant’s competitive fitness *f*_*ij*_ given its local competitors and its infection burden. Hence, the full likelihood of the observed seed output *S* given competitor density is

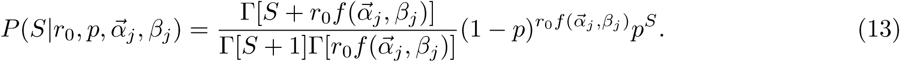

### Estimating competition and foliar infection parameters

We wrote custom MCMC software in Python to perform parameter estimation under our model of competition and infection within a growing season. We suppose that the data correspond to a vector of observed seed outputs, competitor densities, and leaf-area infected for each plant, and we seek to infer each of the *α* and *β* parameters with this method. Our software implements a standard Metropolis-Hastings algorithm. We compute the likelihood function with eqn. 13, and apply an uninformative Γ-shaped prior on *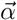* and *β* such that these parameters are constrained to be non-negative. We thinned the observed MCMC traces such that we do not observe strong autocorrelations. We performed 20 independent MCMC runs for each focal species, and found nearly identical parameter estimates across independent runs. Our software is freely available by request and will be posted on the web at a future date.

To assess the performance of our MCMC-based inferences, we performed simulations of seed output under our model and assessed our ability to recapture the parameters that were used to generate the simulated data. For each simulation, we selected a focal plant species *j* and randomly selected competition parameters *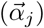* and infection parameters (*β*_*j*_) from uniform priors. We then selected the *p* and *r*_0_ parameters such that the observed mean seed output would exactly match the expected seed output in the simulations. For each plant of species *j* in our dataset, we then simulated a new seed output based on the true density of competitors for this plant and the randomly sampled competition and infection parameters. We then ran our MCMC pipeline on the simulated data, and compared the mean of the posterior distribution of inferred parameter estimates to the true parameter estimates. We find that our method is approximately unbiased and reasonably accurate. A subset of the results of this experiment are plotted in Fig. S5.

Having obtained estimates for the full suite of parameters of our model, we sought to compare predicted seed densities from our model to field observations. For each focal species, we obtained field estimates of the seed density when grown in monoculture and compared them to simulation-based predictions under our demographic model. For each species *j*, we selected a random estimate for the competition parameters *α*_*jj*_ and infection parameters *β* as well as estimates of the other relevant parameters by sampling from the inferred posterior distribution of each parameter and then used a forward simulation under our population dynamic model to obtain an estimate of the final monoculture density. We then compared this set of predicted monoculture densities to the median estimates obtained from our field observations. While monoculture density estimates from our model are noisy, they are concordant with and of similar magnitude to the field estimates (Fig. S6).

### Sensitivity analyses

To better understand which demographic parameters are most important for determining the outcome of competition (*i.e.*, GRWR) for each species, we independently increased each parameter by 5% for each posterior estimate. For simplicity, for each species, we limited this experiment to the set of parameters that directly impact its growth (*i.e.*, its own demographic rates: for each species *j*, only parameters subscripted with *j*). To further reduce the number of parameters, we simultaneously increased all parameters pertaining to the impact of competition on perennial transition from seedling to adult (*i.e.*, *α*_*a*_, *α*_*n*_, and *α*_*s*_), which we then denote as a composite parameter *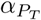*. We included both BFOD infection and fungal foliar infection for this analysis, although the impact of foliar infection was negligible. We compared the predicted GRWR for each species with and without the 5% increase and computed the change in GRWR as ΔGRWR. We report the median ΔGRWR over all posterior samples.

The population growth of both perennial species was most sensitive to the over-summer survival of adult individuals (*ξ*), and essentially insensitive to all other parameters (Fig. S8). *A. barbata* was highly sensitive to *γ* (*i.e.*, the proportion of seeds escaping infection by BFOD), which is unsurprising given the large impact of BFOD on *A. barbata* in the previous analyses and the large burden of infection on *A. barbata* seeds (Fig. S2). *B. diandrus* and *B. hordeaceus* were modestly positively impacted by increasing the probability of establishment for both infected and uninfected seedlings (*ϕ*^*i*^ and *ϕ*^*u*^), and per-capita seed production, *λ*, but negatively impacted by increases in *γ* and interspecific competition with each other.

---

**Figure S1:**
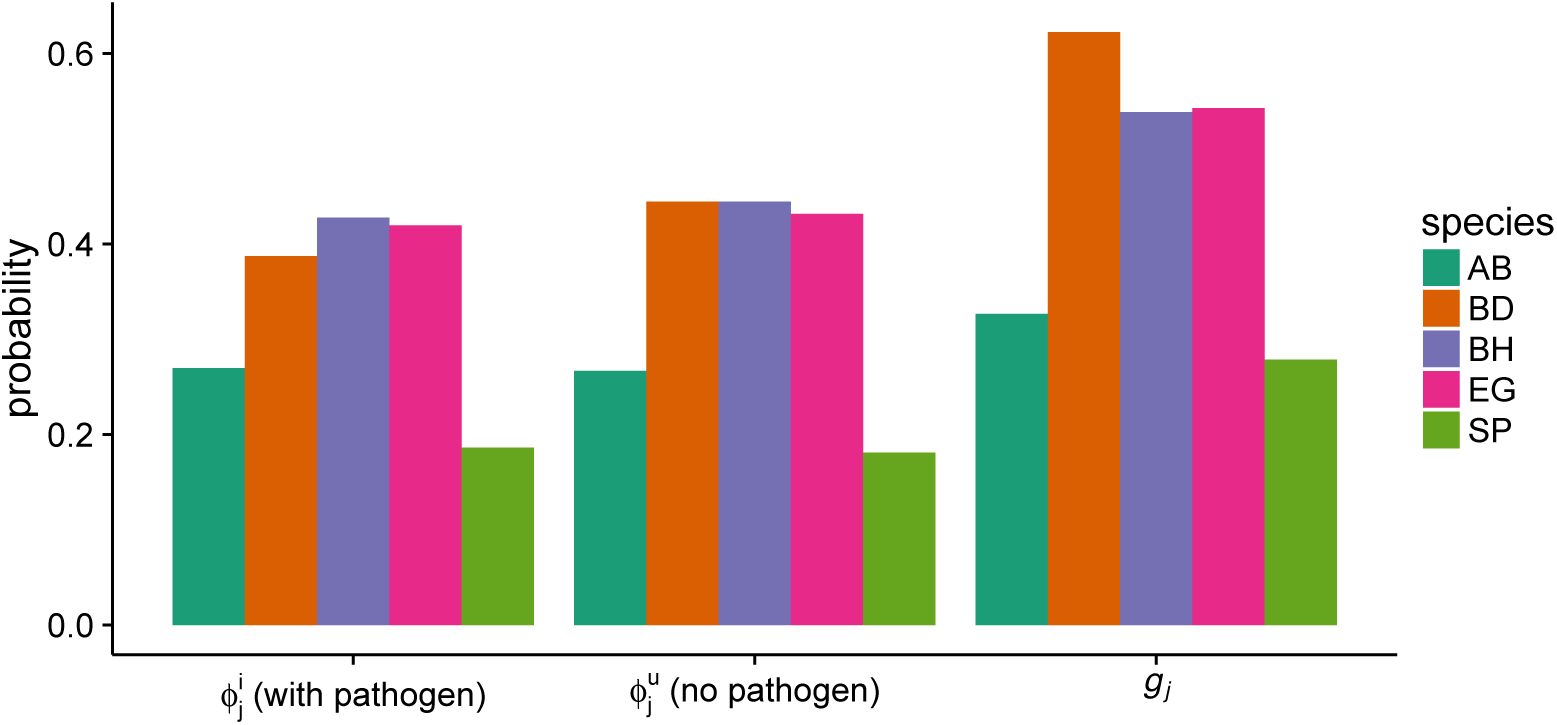
Estimated germination (*g*_*j*_) and establishment (*ϕ*_*j*_) probabilities for each species *j*. Establishment probability was estimated in the presence and absence of foliar fungal pathogens.

**Figure S2:**
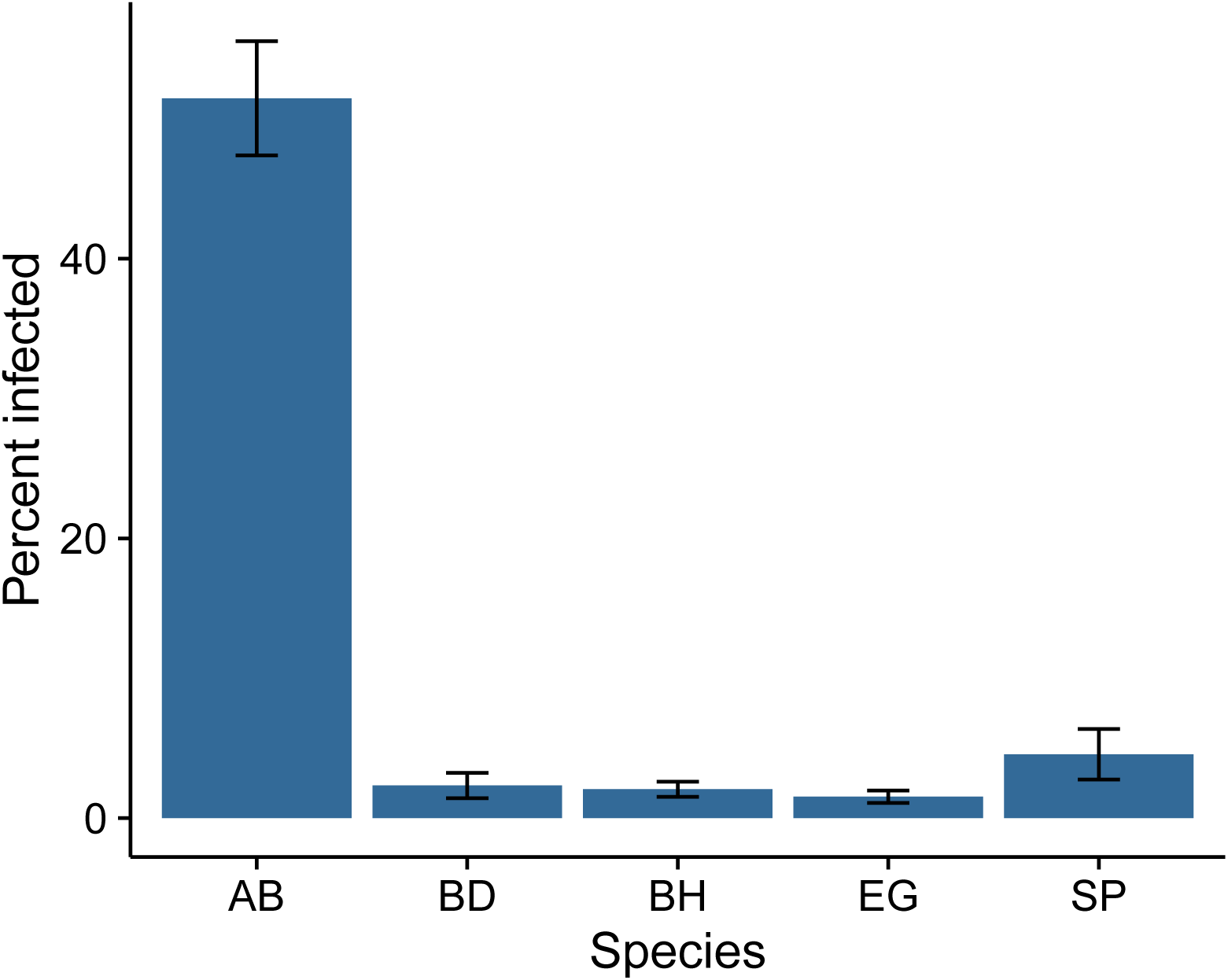
The proportion of seeds that were infected by the Black Fingers of Death pathogen in a seed bag experiment, by species. (Abbreviations: *S. pulchra* (SP), *E. glaucus* (EG), *A. barbata* (AB), *B. hordeaceus* (BH), and *B. diandrus* (BD)).

**Figure S3:**
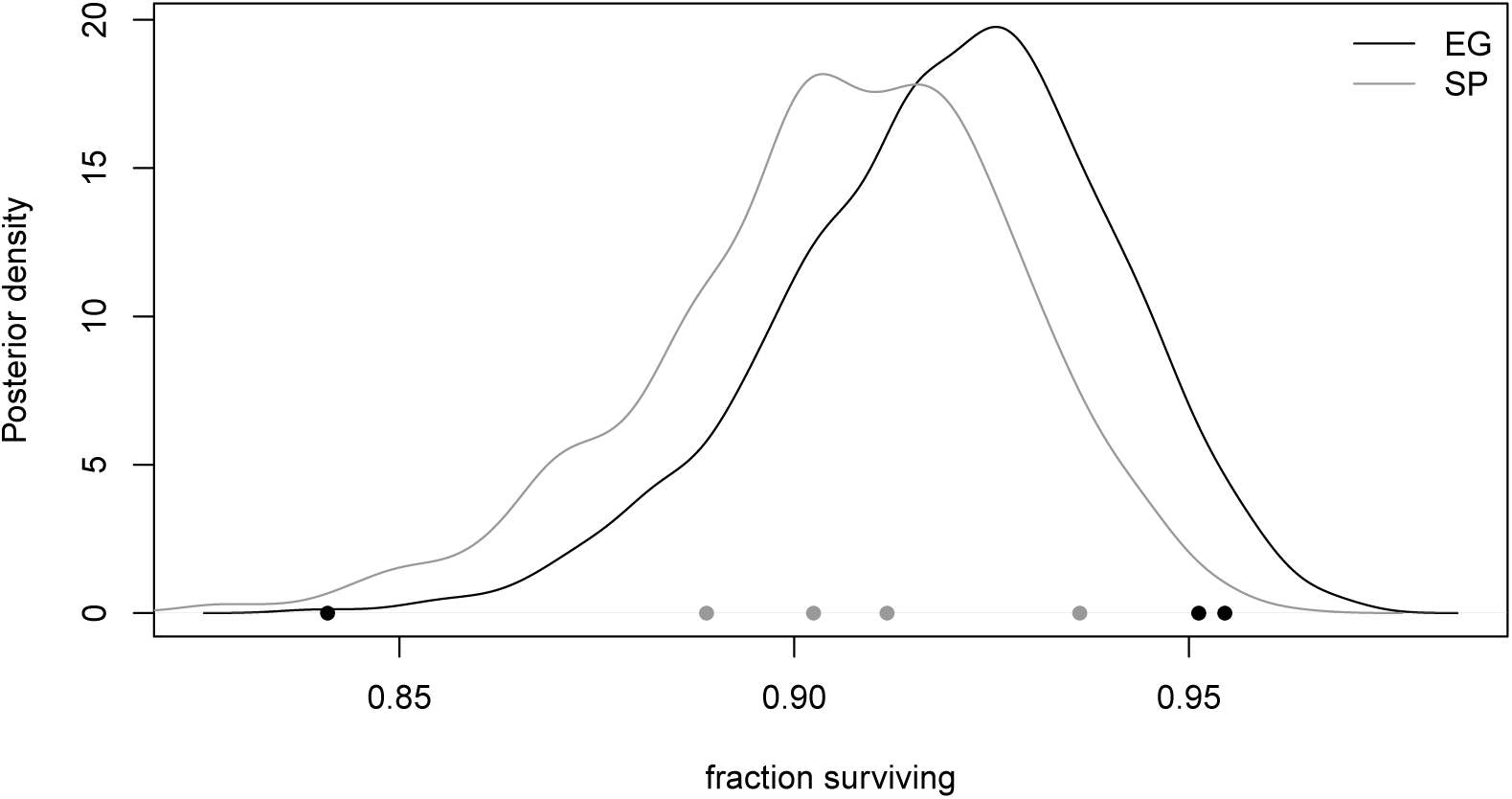
Estimates of the probability of adult perennial over-summer survival rate (which we term *ξ*_*j*_ in our model). The observed fraction of plants surviving across four sampling sites are plotted as points along the x-axis, while the curves represent the posterior density. (Abbreviations: *S. pulchra* (SP), *E. glaucus* (EG)).

**Figure S4:**
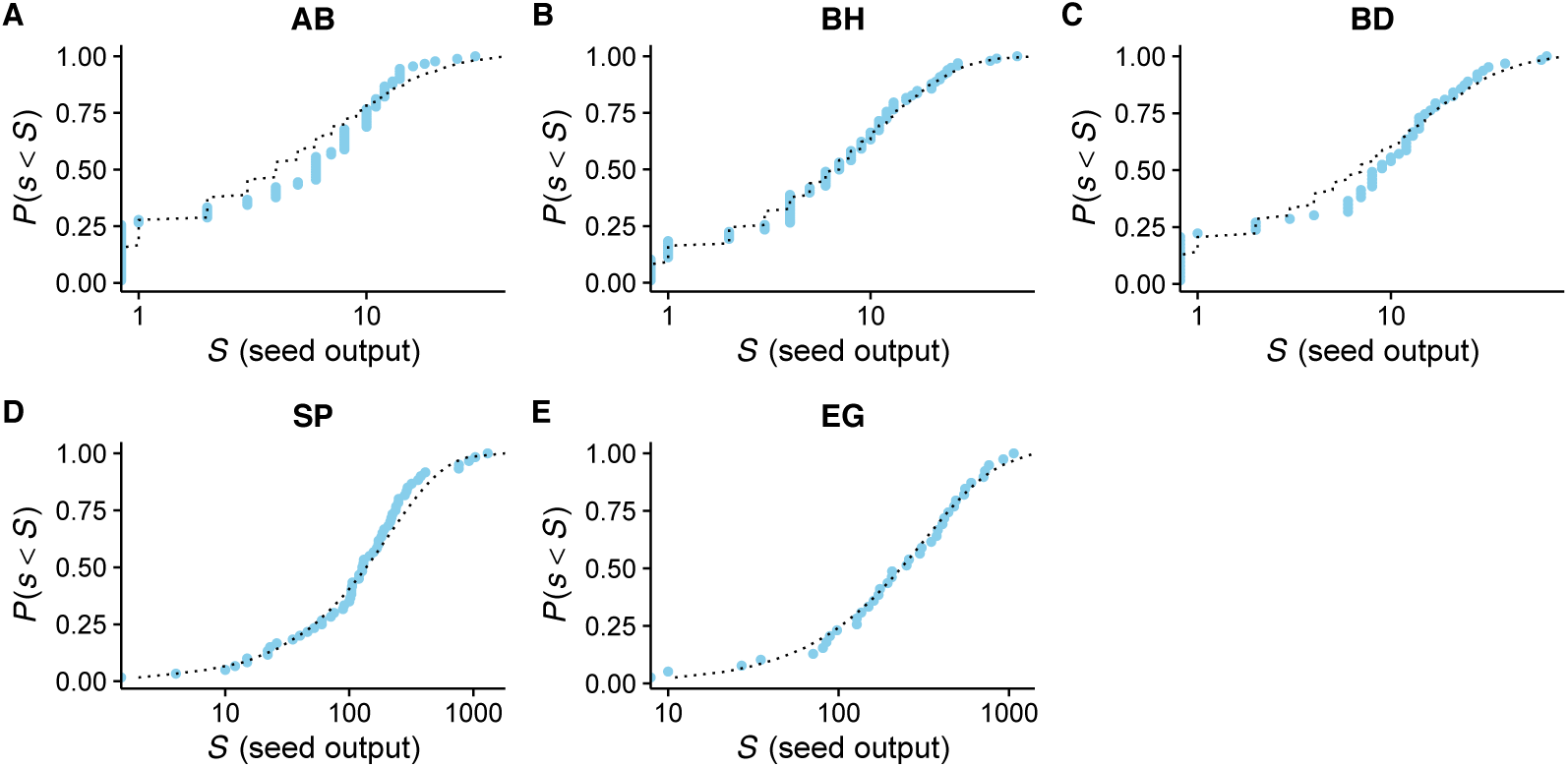
The cumulative distribution of observed seed counts (*P* (*s > S*)) is plotted as a function of seed count *S*. Each point represents the fraction of plants that have a seed count less than *S*. The dashed blue lines show the cumulative distribution of the best fitting negative binomial distribution, obtained my maximum likelihood. (Abbreviations: *S. pulchra* (SP), *E. glaucus* (EG), *A. barbata* (AB), *B. hordeaceus* (BH), and *B. diandrus* (BD)).

**Figure S5:**
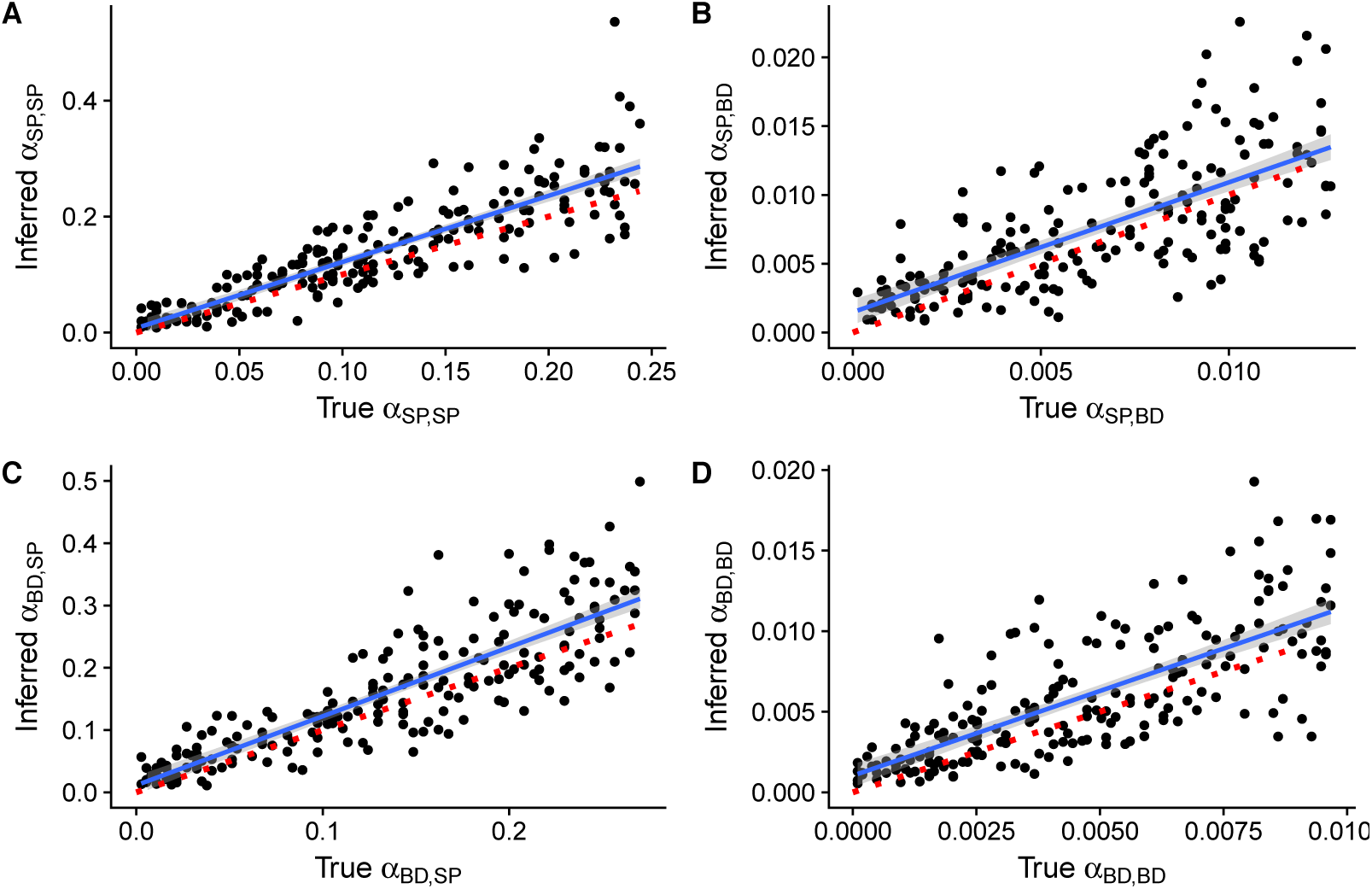
True and inferred parameter values as determined by our MCMC method, as obtained from simulation. The inferred values represent the mean of the inferred posterior distribution. The lines and envelopes (in gray) were fit using a simple linear model of the form *y mx* + *b* with the method `"lm"` in the function `geom_smooth` in `ggplot2`, while the red dashed lines show the diagonal (*i.e.*, the expectation if inference were perfect). *α*_*BD,SP*_ is the impact of *S. pulchra* on *B. diandrus*. (Abbreviations: *S. pulchra* (SP) and *B. diandrus* (BD)).

**Figure S6:**
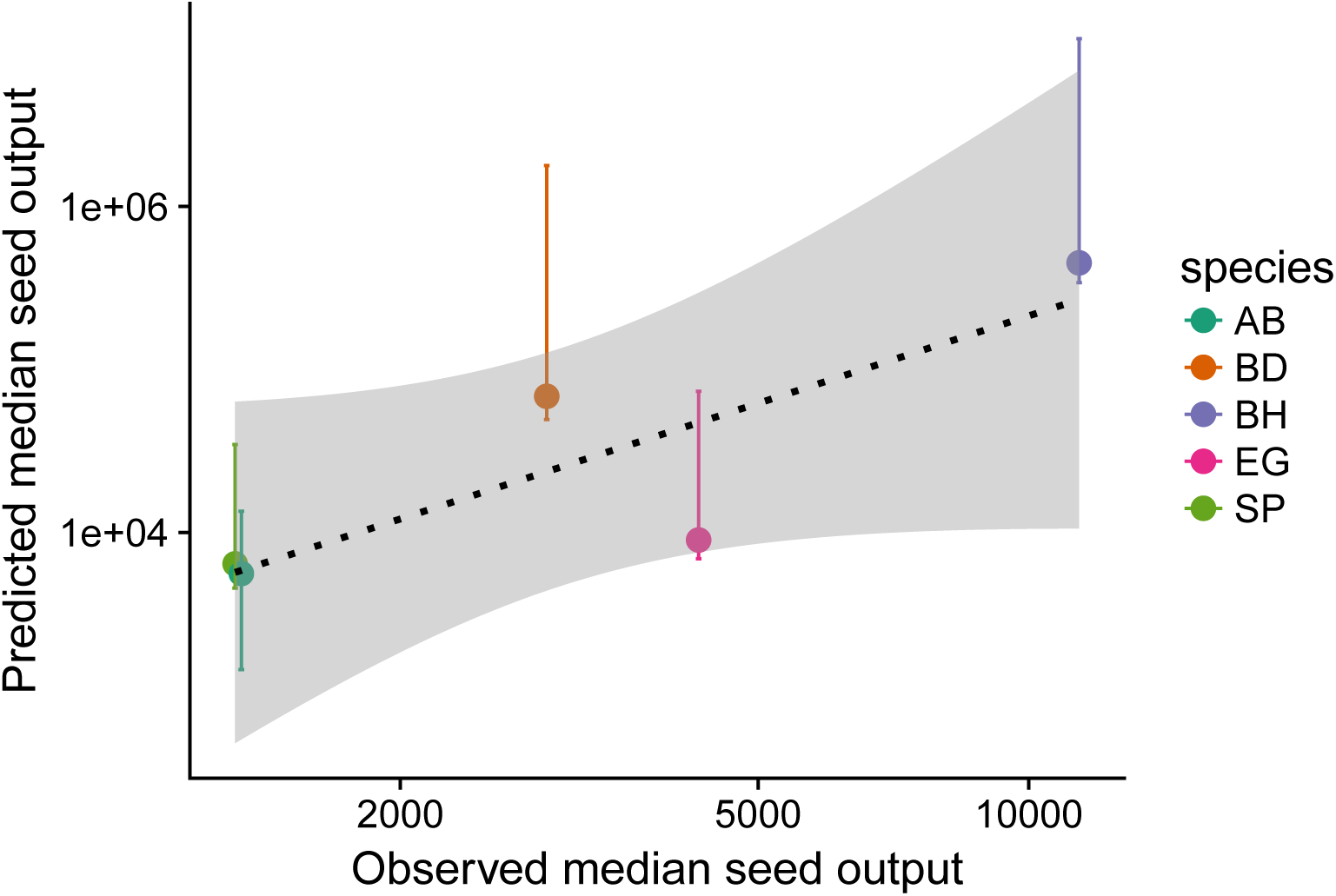
Observed median monoculture seed output densities plotted against those predicted by our population dynamic model. Error bars represent the inner 95% of parameter estimates from our model. The line and envelope (in gray) were fit using a simple linear model of the form *y mx*+*b* with the method `"lm"` in the function `geom_smooth` in `ggplot2`. (Abbreviations: *S. pulchra* (SP), *E. glaucus* (EG), *A. barbata* (AB), *B. hordeaceus* (BH), and *B. diandrus* (BD)).

**Figure S7:**
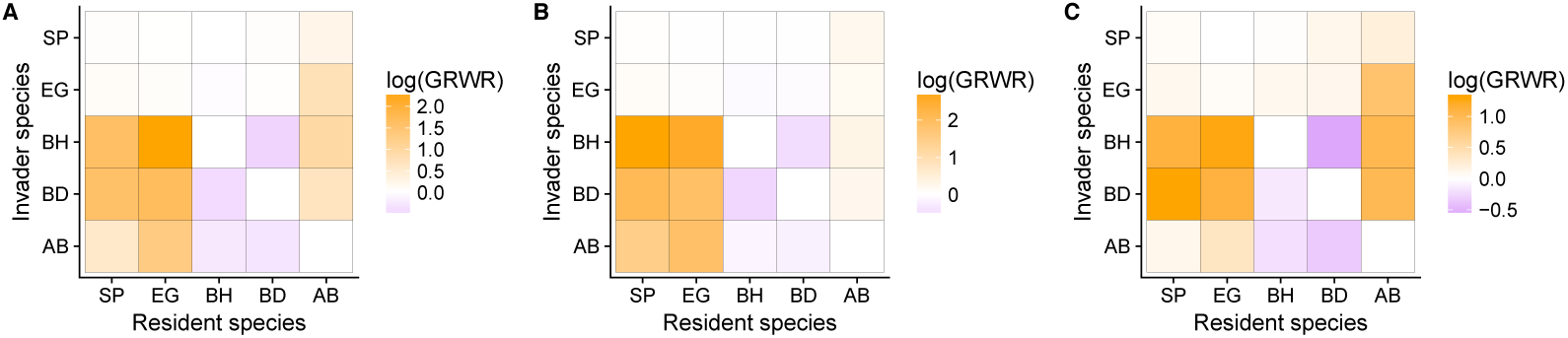
Comparison of pairwise invasion analyses using A: all data (competition plots and transects), B: only the data from the competition experiment, or C: only data from the transects. (Abbreviations: *S. pulchra* (SP), *E. glaucus* (EG), *A. barbata* (AB), *B. hordeaceus* (BH), and *B. diandrus* (BD)).

**Figure S8:**
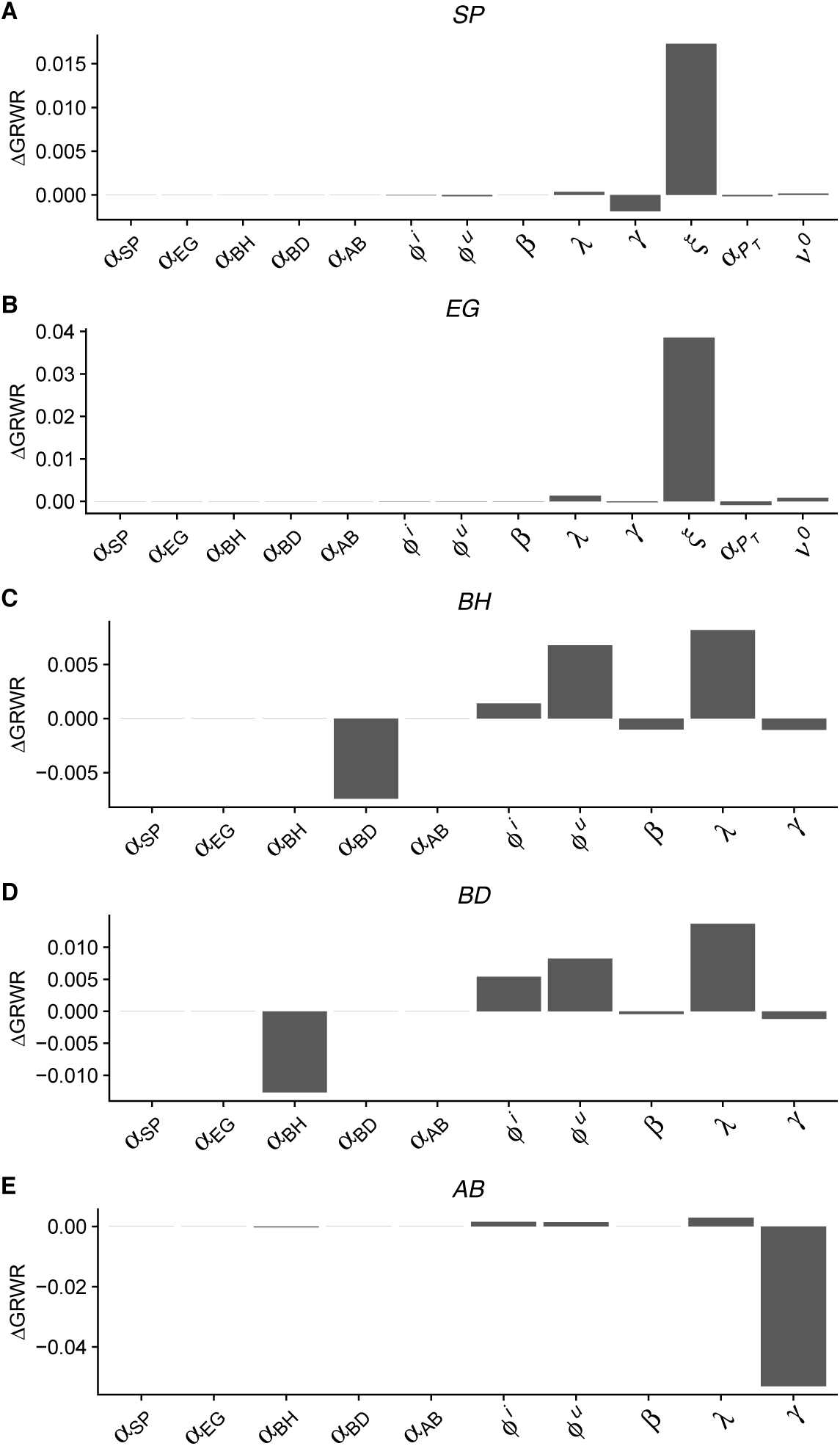
Sensitivity analysis results. For each species, we plot the proportional difference in GRWR when the corresponding parameter is increased by 5%. *α*_*SP*_ denotes the impact of *S. pulchra* on the focal species, *α*_*EG*_ represents the impact of *E. glaucus* on the focal species, etc. Parameters are defined in Table 1. (Abbreviations: perennial native species (*S. pulchra* (SP), *E. glaucus* (EG)), annual exotic species (*A. barbata* (AB), *B. hordeaceus* (BH), and *B. diandrus* (BD))).

